# Host stress hormones affect host, but not vector, infectiousness for West Nile virus

**DOI:** 10.1101/2021.05.20.444978

**Authors:** Lynn B. Martin, Meredith E. Kernbach, Kyle Koller, Nathan D. Burkett-Cadena, Thomas R. Unnasch

## Abstract

Hormones that help hosts cope with stressors also affect how hosts regulate the processes that influence their susceptibility to parasites as well as their propensity to transmit pathogens to other hosts and vectors. In birds, corticosterone (CORT), influences timing of activity, feeding behaviors, and various immune defenses that influence the number and outcomes of host interactions with vectors and parasites. No study to our knowledge, though, has investigated whether CORT in hosts affects the extrinsic incubation period (EIP) of a vector for a virus, one of the strongest drivers of vector-borne disease cycles. Our goal here was to discern whether experimental CORT alterations in zebra finches (*Taeniopygia guttata*) affected EIP for West Nile virus (WNV) in the mosquito, *Culex quinquefasciatus*, a common vector of WNV and other infections in the southern US. We experimentally manipulated CORT in birds, infected them with WNV, and then investigated whether EIP differed between vectors fed on CORT-treated or control birds. Although CORT enhanced WNV viremia in hosts, as we have observed previously, we found no effects of CORT on vector EIP or post-feeding mortality rates, another important component of epidemiological models. These results, plus our prior observations that CORT enhances host attractiveness, indicate that some but not all stages of host-vector-virus interactions are sensitive to host stress.

## Introduction

Stressors, both anthropogenic and natural, affect the outcomes of infectious diseases at multiple levels of biological organization (Becker et al. 2019; Martin et al. 2016). For individuals, stressors can alter the behaviors that influence exposure probability to infected conspecifics and vectors (Barron et al. 2015). They can likewise affect the immune system processes that determine how effectively parasites are controlled (Dhabhar 2009) as well as the degree to which infected hosts experience morbidity and mortality (i.e., sickness behaviors and pathology) (Adelman and Martin 2009). Individual-level effects of stressors can also scale up to influence the emergence, persistence, and rates of spread of various infections in populations (Altizer et al. 2018; Hawley and Altizer 2011; Martin et al. 2019a). Whereas the underlying mechanisms of such outcomes are numerous, inter-individual variation in host-parasite interactions can impinge on community level disease dynamics (Plowright et al. 2017).

For vectored infections, particular stages of host-parasite interactions have gained little to no attention with respect to stressors. One is the extrinsic incubation period (EIP) (Richards et al. 2007), the rate at which vectors that rely on host blood for viability and especially reproduction become infectious from infected host blood-meals. Epidemiologically, EIP is a strong influence on disease prevalence and emergence (Foppa and Spielman 2007). Nevertheless, for most infections, we know little about whether stressors in hosts affect it, much less by what mechanisms (Dhondt and Dobson 2017). This paucity of study is somewhat surprising given that host bloodmeal composition can vary extensively among individuals in response to stressors, which could affect the viability of internal parasites prior to vector infection as well as the propensity of parasites to infect vectors (Hurd et al. 1995). Indeed, glucocorticoid hormones, which are commonly altered when vertebrates are exposed to unexpected or enduringly challenging stressors (Romero and Wingfield 2015), can alter host blood characteristics in such a manner that vector infection probability is changed (Beck et al. 2016). Likewise, glucocorticoids can affect host defensive behaviors towards vectors (Gervasi et al. 2016), which could alter the bloodmeal size taken by foraging vectors.

In the present study, our goal was to query directly whether glucocorticoids in avian hosts affected vector responses to a pathogen, West Nile virus. We did not expect direct effects of CORT on vector traits, as vectors lack glucocorticoid receptors. We expected any CORT effects on EIP and mortality to arise indirectly, via the quantity or quality of host blood meals. In some vectors, larger blood meals can result in larger vector clutch sizes (Prasad 1987), probably because large blood meals contain more protein for vitellogenesis (Briegel 1990). Blood meal composition was also a plausible mechanism for any effects of CORT on vector EIP for WNV (Shieh and Rossignol 1992); chronic CORT elevation in many vertebrates can cause hyperglycemia and alter lipid levels (Dallman and Bhatnagar 2001) and decrease hematocrit (Beck et al. 2016; Gervasi et al. 2016), and most of the protein in blood is concentrated in hemoglobin (Hurd et al. 1995). Although such effects are not observed in all avian species in response to all stressors (Cyr et al. 2007), CORT-mediated alterations to blood composition could plausibly affect vector EIP (Vaidyanathan et al. 2008). We did not test these pathways by measuring bloodmeal size and composition, as efforts to do so would have complicated study design and potentially masked CORT effects on vector traits (due to excessive handling of birds). Here, our main focus was to test whether avian corticosterone (CORT) affects vector EIP in an important arboviral system, the interaction among West Nile virus (WNV), one passerine host, the zebra finch (*Taeniopygia guttata*), and one of the most common vectors of WNV in the southeastern US, *Culex quinquefasciatus* (Rochlin et al. 2019).

Our choice of this system was a natural extension and progression of our previous findings that experimental corticosterone (CORT) manipulation made individual zebra finches twice as attractive (Gervasi et al. 2016) and more infectious (Gervasi et al. 2017a) to vectors, which probably equates to higher competence in natural systems. Indeed, birds that attract more vectors and also circulate virus for long periods above thresholds where vectors are likely to become infectious themselves should generate more infectious, and thus be more competent, than those that clear virus quickly or never let it reach transmissible titers. Here, we sought to determine if CORT might further enhance host competence by shortening vector EIP; such effects when coupled with greater attractiveness and infectiousness of hosts could make host stress a very strong driver of disease epidemics. As in our previous work, we implanted finches with CORT (or sham controls), allowed CORT to change in circulation for several days, exposed finches to WNV experimentally, allowed mosquitoes to feed on birds at peak viremia (4d post-infection), then queried whether CORT treatment affected mosquito mortality rate and EIP, specifically the rate at which WNV reached mosquito salivary glands.

## Methods

### Study organisms

We studied WNV in zebra finches and *Cx. quinquefasciatus* for three reasons. First, we sought to complete a series of studies on the same host, vector, and virus interactions for direct comparisons so that we could ultimately examine simultaneously all the pathways by which host stress hormones might affect local disease dynamics. As above, we had studied CORT effects on vector feeding choice, anti-vector behaviors, host and resistance and tolerance of WNV previously (Gervasi et al. 2017a; Gervasi et al. 2016). EIP was a logical and important next step in the epidemiological pathway. Second, we chose *Cx. quinquefasciatus* because it is one of the more common mosquito vectors of WNV in the southeastern US (Burkett-Cadena 2013) and was also the focus of our prior work (Gervasi et al. 2016). Zebra finch responses to WNV, too, had been studied before by us (Gervasi et al. 2017a) and others (Hofmeister et al. 2017; Newhouse et al. 2017). Our original choice of this avian species was due to its sequenced genome, simple husbandry, and the ability to breed it in captivity so as to obtain large sample sizes. Captive breeding also ensured no prior exposure, ecologically or evolutionarily, to WNV. Third, we chose WNV because it is the most broadly distributed arbovirus and most important causative agent of viral encephalitis worldwide (Paz 2019). Since its introduction to the US in 1999, WNV has infected almost 40,000 humans, 1,667 of whom died from the neuroinvasive form (WNND). WNV is predominantly an infection of passerines and ornithophilic vectors, and continues to have large consequences for passerines (LaDeau et al. 2007), although it can be transmitted by as many as 45 vector species (Kramer et al. 2007; Marra et al. 2004).

### Bird husbandry and CORT and WNV exposure

We obtained 18 adult zebra finches from an active breeding colony maintained at the University of South Florida. Specific birds used in the study came from groups of 15 - 18 birds housed together in free-flight cages (90×60×60 cm). Birds were randomly chosen from the above groups and assigned to one of three treatments: control (sham, CORT+, or CORT++ (Ouyang et al. 2013)). During the experiment itself, birds were housed singly in 30.48 cm^3^ mosquito-proof cages (BioQuip, Rancho Dominguez, CA, USA, product # 1450 BSV), but a clear plastic panel on one side of each cage enabled birds to remain in sight of each other and a mesh covering on another side of each cage permitted audial contact of conspecifics. Birds were individually housed for 3 days prior to hormone implantation, allowed to recover from surgery for 2 days, then moved to an Animal Biosafety Level (ABSL) 3 facility where they acclimated for another 24 h before being exposed to WNV. For the duration of the study, all birds received ABBA 1900 exotic finch food (ABBA Products Corp., Hillside, NJ), photoperiod was kept at 13h light:11h dark (on at 0600 and off at 1900), and room temperature and relative humidity were maintained at ~21°C and ~50%, respectively. All birds were housed in proximity to each other for the study duration, and all procedures complied with approved USF animal care and use and biosafety protocols.

For CORT treatments, we implanted 18 birds total subcutaneously (s.q.), some with CORT (n = 12) and some (n = 6) with sham treatments (7mm long; inner diameter 1.5 mm, Dow Corning, Midland MI, product #508-006); importantly, though, we implanted 6 birds with 2 CORT-filled silastic tubules (3 males and 3 females) and 6 birds with 1 CORT-filled tubules (3 males and 3 females). Control birds (3 males and 3 females) received an empty silastic tubule (Gervasi et al. 2017b; Gervasi et al. 2016; Ouyang et al. 2013). Implantation of different numbers of tubules was intended to cause dose-dependent elevations of CORT in the blood. All tubules were sealed (Dow Corning, Midland, MI, product #732) several days prior to implant, but minutes before each implantation, a 0.5 mm hole was bored through each implant to optimize efflux of hormone (Ouyang et al., 2013). Tubules were then implanted on one flank of each bird while it was sedated with light isoflurane anesthesia. After implantation, wounds were sealed with surgical adhesive (Vetbond, 3M, St. Paul, MN, product #1469). All birds returned to normal activity (perching and feeding) within minutes of implantation.

Once CORT had time to take effect (3 days after implantation), each bird was exposed to 1×10^7^ PFU WNV (NY99) via s.q. injection (Gervasi et al., 2017). Blood samples (75ul) were then taken around 0800h 4 days after WNV exposure (d.p.e.) to quantify viremia in birds; after blood sampling, serum was removed from samples and stored at −40°C until RNA extraction. On day 4 post WNV exposure, we introduced 23 mosquitoes into the cage of each bird 1h before lights-out (~1900h). We allowed mosquitoes to feed on birds until the following morning (0600), at which point several blood-fed mosquitoes (based on visual inspection of abdomens) were aspirated from each bird cage into separate plastic containers (Glad, 32 oz bowls with 1.5 oz water-filled plastic cups in the bottom; all mosquitoes housed singly) for the next 12 days. Plastic domiciles were also lined with wet paper towels to foster high humidity as well as a paper card laden with honey for collection of mosquito saliva from which we could later assess WNV presence (Burkett-Cadena et al. 2016). Fresh ‘honey-cards’ were added to each mosquito domicile every other day, and used cards were removed and stored (−80°C) until extractions for WNV detection. We also monitored mosquito survival over this same period. Uninfected birds were not included in this study for two reasons: i) we had insufficient space in the ABSL-3 facility for additional birds, and ii) our goal was to assess CORT effects on vector EIP, which requires WNV infection.

### Mosquito husbandry

A laboratory colony of *Cx. quinquefasciatus* was established using a previous colony (generation > F100) from Indian River County, FL. Larvae were reared at 28°C and maintained under a 14:10 (light:dark) cycle. Three to four egg rafts (200-300 eggs each) were placed in larval rearing pans (45.7 cm × 53.3 cm × 7.62 cm) containing approximately 3L tap water. Larvae were fed daily with (20 mg/mL) of 1:1 Brewer’s yeast and lactalbumin. Pupae were transferred to containers with ~250 mL of clean water and placed into cages (30.48 cm^3^) for emergence. Adults were provided 20% sucrose *ad libitum* until 24 hours prior to experiments, when was removed. Females were transported to USF in one-liter cardboard holding containers with mesh screen tops where they were held without food for <2d until introduction to bird cages. Only females of uniform age were used (typically 7-10 days post emergence).

### WNV quantification in avian sera and mosquito saliva

We used quantitative real-time polymerase chain reaction (qRT-PCR) to measure WNV viremia in birds (Gervasi et al., 2017; Burgan et al., 2018). Briefly, viral RNA was extracted from serum samples with the QIAmp Viral RNA kit (Qiagen Cat. No. 52906). We used 10 μl of serum diluted in 130 μl sterile PBS and followed the kit protocol for all steps. We then used a one-step Taq-based kit to quantify viral RNA in samples (iTaq Universal Probes One-Step Kit; Bio-Rad Cat. No. 1725141). As in the past, our forward primer sequence was 5’ CAGACCACGCTACGGCG 3’; reverse sequence was 5’ CTAGGGCCGCGTGGG 3’; and our WNV probe sequence was 5’ [6~FAM] CTGCGGAGAGTGCAGTCTGCGAT [BHQ1a~6FAM]. All samples were measured in duplicate, and a negative control and a known WNV-positive control were run concurrently on all plates. WNV standard curves were generated from serial dilutions of the same stock virus used in finch inoculations, in which viral titer was quantified previously using Vero cell plaque assay. All samples on all plates were captured by our standard curves.

### WNV quantification in mosquitoes

WNV positivity in mosquito saliva (i.e., on honey cards) was assessed at 6, 8, 10− and 12-days post-feeding on birds, as viral dissemination and maturation typically takes about 10 days in *Cx. quinquefasciatus* under ideal conditions (Richards et al. 2007). We queried infections status at all 4 time points to ensure that any expediting or slowing effects of CORT were captured.

### Data analysis

We used ANOVAs to compare effects of CORT treatments on avian viremia, the number of mosquitoes surviving the overnight feed, and the number becoming blood-engorged during that feed; Levene’s test indicated no heteroscedasticity among groups, and data were normally distributed. We used Cox survival analyses to assess host CORT treatment and viremia effects on mosquito mortality and infection rates. We used SPSS v24 for all analyses, and GraphPad Prism v6 for all figures, setting alpha to 0.05.

## Results

### Hormone implant effects on avian viremia

CORT effects on viremia were not statistically significant when we compared all three groups against each other (F_2, 16_ = 4.5, P = 0.09, Fig. 1A), but the tendency for CORT-treated birds to have higher viremia and the precedent from our prior work that our two implant categories rarely differed with respect to effects on finch WNV responses motivated us to combine the two CORT groups here into a single group (Gervasi et al. 2017a; Gervasi et al. 2016). That approach revealed that viremia was higher in CORT-implanted birds compared to controls (F_1,16_ = 5.9, P = 0.03; Fig. 1B), as observed in the past. Sex of birds did not influence viremia (F_1, 16_ = 0.49, P = 0.50).

**Fig. 1.**
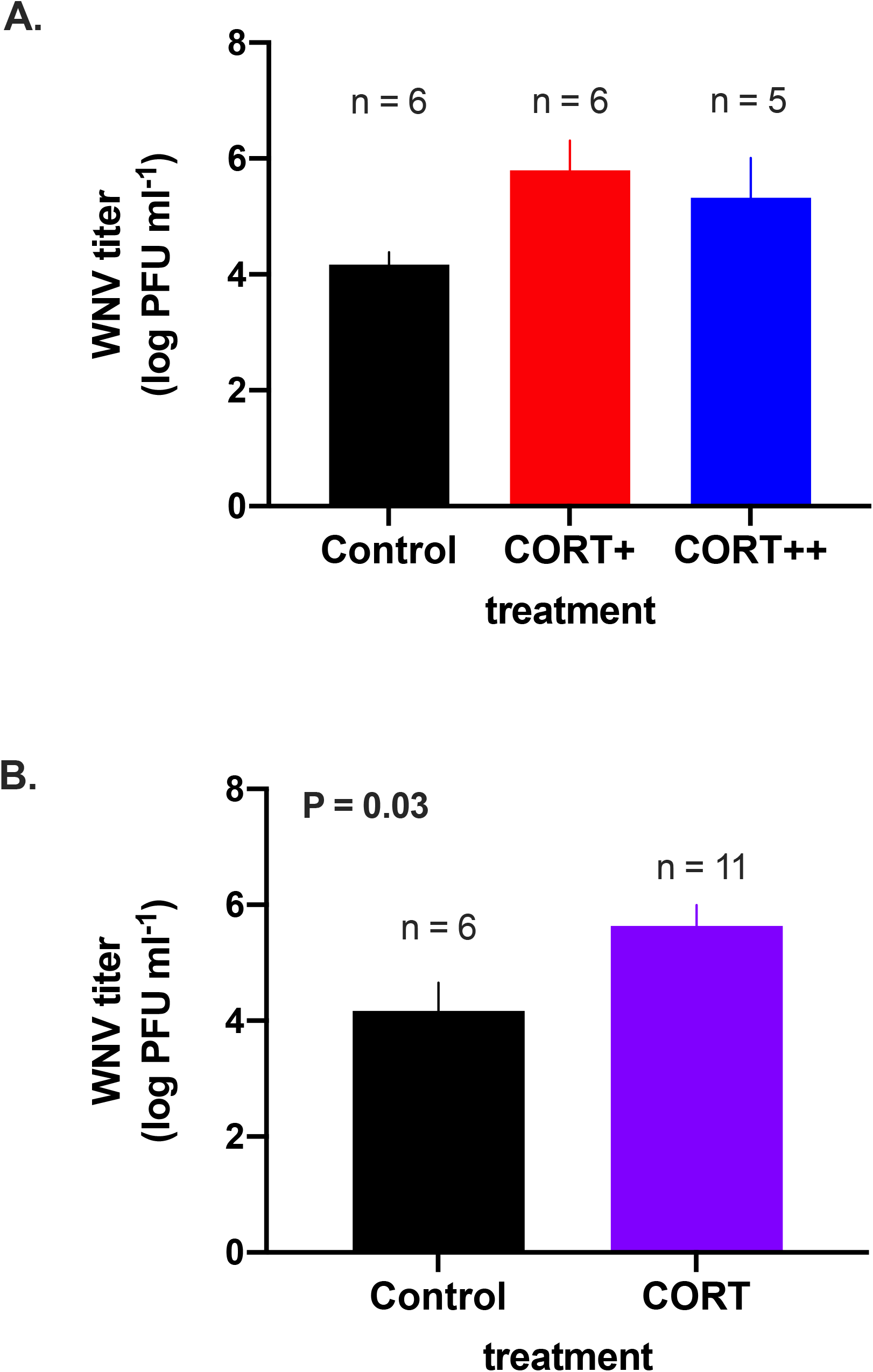
Effects of corticosterone (CORT) on zebra finch responses to West Nile virus infection 4 days post-exposure. A) viremia did not differ among groups when CORT-treatment groups were compared separately to controls, but B) when CORT-treatments were collapsed into one group, treated birds had higher viremia than controls. Bars depict means +/−1 SE.

### CORT implant effects on mosquito overnight survival and feeding success

CORT treatment did not affect the number of mosquitoes remaining alive after they were co-housed with birds overnight (F_2,16_ = 2.61, P = 0.11; Fig. 2A), nor the number of fully-engorged mosquitoes collected from bird cages the morning after trials (F_2,16_ = 1.86, P = 0.19; Fig. 2B). When we merged CORT+ and CORT++ birds into a single ‘CORT’ group, we observed similar non-significant outcomes (mosquitoes alive: F_1,16_ = 0.65, P = 0.43; blood-engorged mosquitoes recovered: F_1,16_ = 0.93, P = 0.35). However, whereas for each control and CORT+ bird, at least some mosquitoes fed successfully, blood-fed mosquitoes were collected from only 3 CORT++ birds. One male CORT++ bird died prior to mosquito exposure, and cages of 2 CORT++ birds contained no blood-fed mosquitoes the morning of collection.

**Fig. 2.**
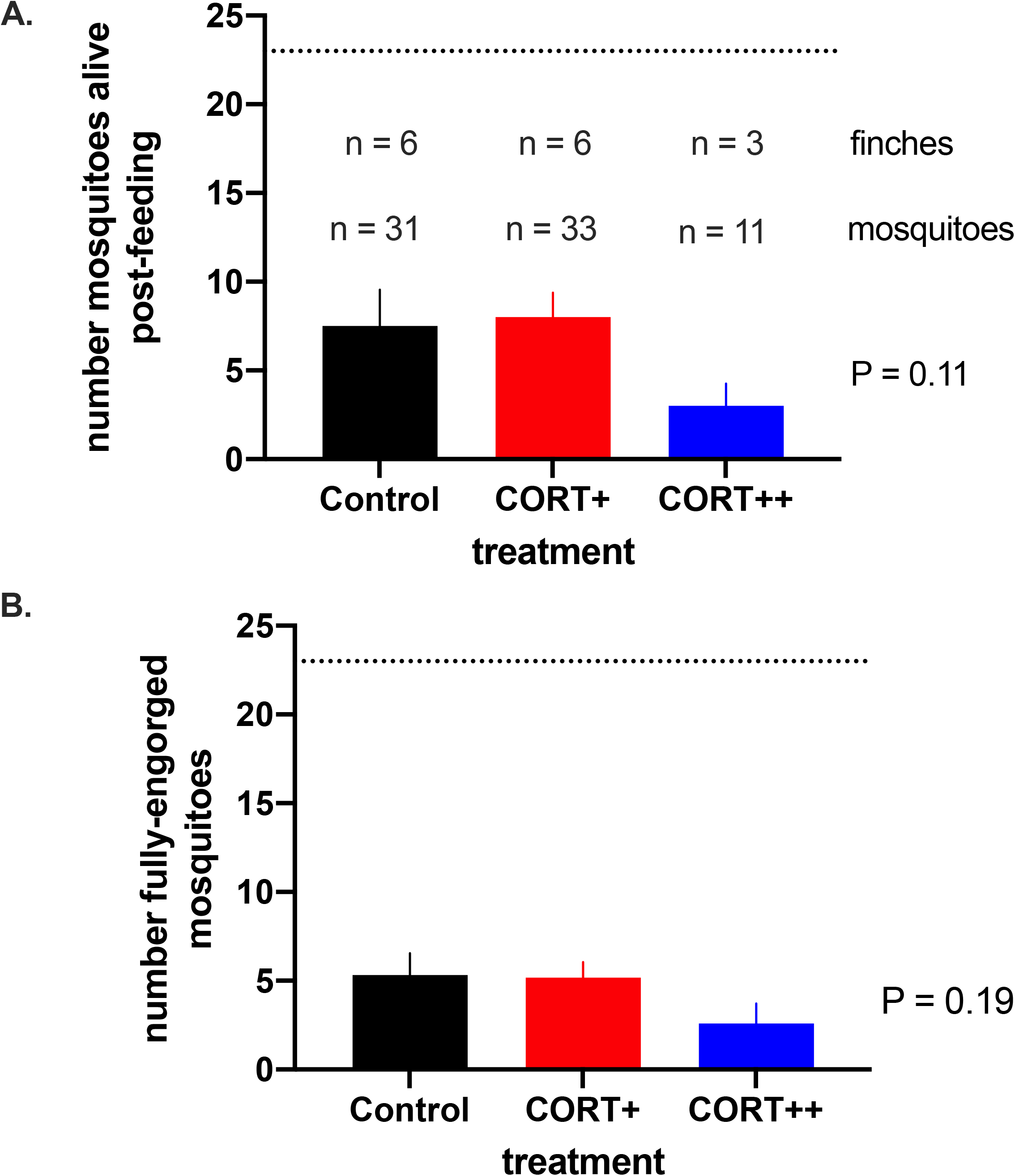
Corticosterone treatment did not affect A.) number of mosquitoes found alive or B.) number of fully engorged mosquitoes the morning post-feeding. Bars are means +/− 1SE; dotted line denotes total number of mosquitoes (n = 23) to which birds were exposed the prior evening. In A., numbers above bars denote total birds from which mosquitoes were collected the following morning and total mosquitoes studied for WNV infectivity.

### CORT effects on mosquito mortality rate post blood-feeding

Fifteen mosquitoes died prior to assessing WNV infection status, which explains the disparity in sample sizes between Fig. 3A and 3B. When all three CORT treatments of birds were considered separately, neither CORT treatment, nor viremia in birds, nor their interaction affected mosquito mortality rates (omnibus χ^2^_5_ = 4.2, P = 0.52). Analyzing both CORT treatments as a single group in a similar model produced similar non-significant results (omnibus χ^2^_3_ = 1.1, P = 0.77).

**Fig. 3.**
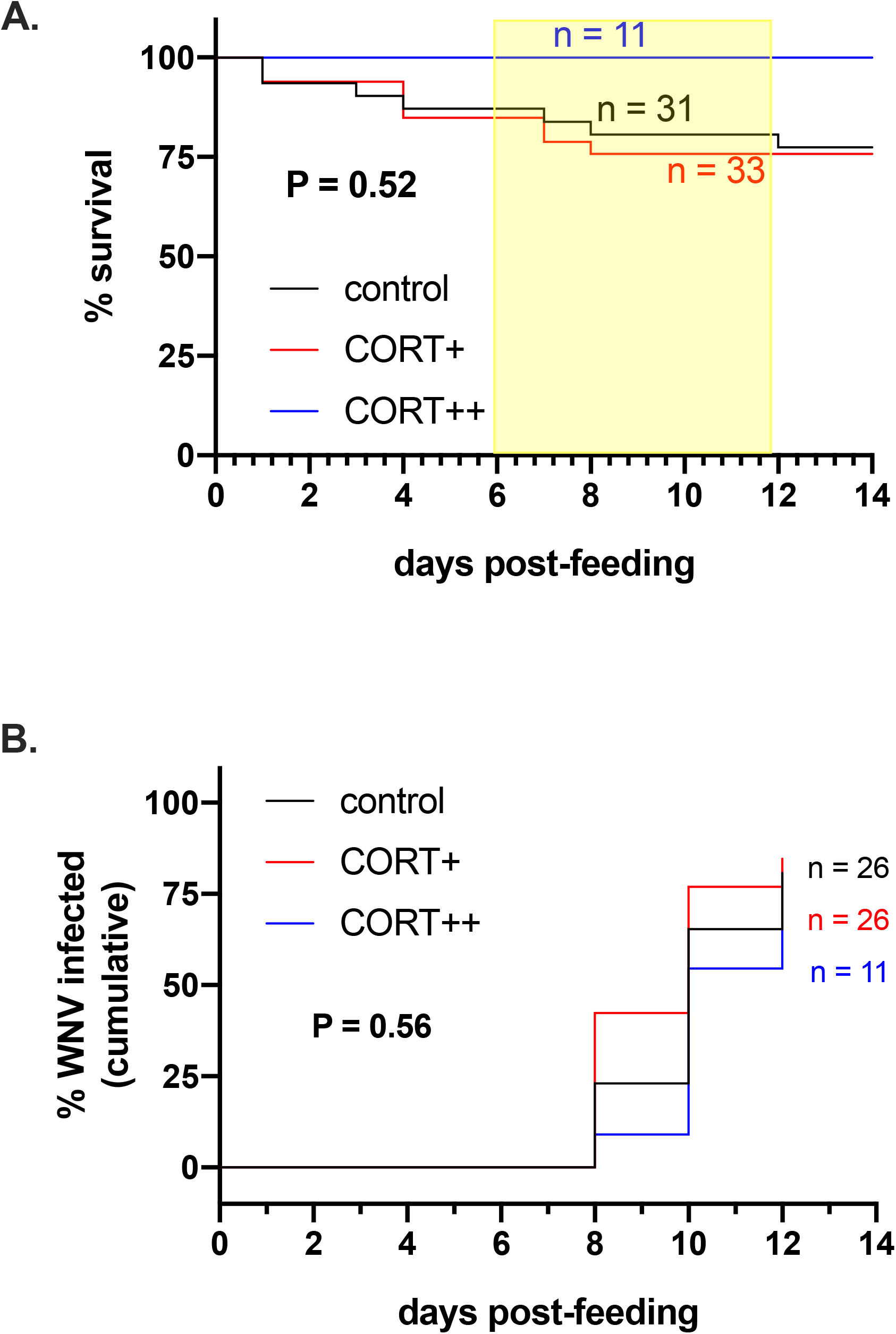
No effects of corticosterone treatment of finches on A. mortality rate or B. WNV infection rate of *C. quinquefasciatus*. Lines denote A. survival or B. cumulative infection curves (% of birds) over the 14-day monitoring period. Shaded area in A. denotes period of screening of mosquito saliva for WNV (i.e., data comprising figure 3B). Sample sizes depicted in color denote mosquitoes in each respective group at the time mortality or WNV infection surveillance began.

### CORT effects on extrinsic incubation period for WNV

Neither CORT, nor viremia in birds nor their interaction affected the rate at which mosquitoes became infected with WNV. Results were consistent when we analyzed the two CORT groups separately (omnibus χ^2^_5_ = 3.9, P = 0.56; Fig. 3B) and when CORT groups were collapsed and compared to controls (omnibus χ^2^_3_ = 3.6, P = 0.30).

## Discussion

As in our previous work, we found that CORT treatment elevated WNV titer in zebra finches, compared to controls (Gervasi et al. 2017a; Gervasi et al. 2016). CORT did not influence the number of mosquitoes that survived a night of feeding on birds, though, nor the number of mosquitoes that successfully obtained a blood-meal from hosts. Likewise, neither CORT treatment nor viremia from the avian host affected mosquito mortality rates or the rate at which WNV became detectable in vector saliva (i.e., EIP). Below we discuss the ramifications of these results, particularly in relation to our other work that CORT affects vector choice of hosts and host viremia.

### CORT effects on avian responses to WNV

We found similar enhancive effects of CORT on WNV viremia in this species. Given the small sample sizes of birds we had to study, enhancive effects of CORT on viremia were only observed when CORT treatment groups were combined. In all previous studies involving this implantation technique in finches exposed to WNV, we have taken an identical approach, combing CORT treatments into a single group. Our original motivation for studying two different implant types was to identify protective levels of CORT (Martin 2009), assuming that the single implant would be protective and the double implant detrimental to host resistance and/or health. However, in no study yet have we been able to detect such subtle effects including another whereby we transiently elevated CORT via injection (Martin et al. 2019b). Injected and implanted CORT elevated circulating CORT to the same levels in birds, but viremia was only increased in implanted birds relative to controls. Injected CORT was not protective, as injected and control bird viremias were indistinguishable.

In previous publications (Gervasi et al., 2016, 2017), we have extensively discussed the potential limitations of our approach to manipulating CORT, a hormone that is tightly regulated by multiple positive and negative feedback loops (Romero and Wingfield 2015). We are aware of the great difficulties of simulating natural fluctuations in circulating concentrations of this hormone (MacDougall-Shackleton et al. 2019), and we do not claim that our method is ideal. Here and elsewhere, our implants were intended to simulate physiological responses to a brief food shortage, prolonged weather event, or comparable stressor, which would be expected to alter CORT concentrations for the same period as our method. Nevertheless, over the same 2-day periods, expression of glucocorticoid receptors as well as other intermediaries of CORT regulation (i.e., binding globulins in blood, cytosolic co-receptors, etc.) are apt to change in response to CORT manipulation (Romero and Wingfield 2015), making variation in CORT concentrations among individuals difficult to interpret functionally. As other methods of CORT administration (e.g., injection, addition to drinking water, osmotic pumps, etc.) also have practical and inferential limitations, our perspective has been to emphasize the inferential limits of our study while also appreciating that we cannot experimentally infect wild birds with WNV after natural stressors, a preferable but ethically impossible design. In the end, we favored experimental tractability and replicability, but we agree that pairing controlled studies such as ours with creative, complementary fieldwork will ultimately be the best option for this complex topic.

### Lack of CORT effects on vector mortality and EIP

CORT treatment did not affect the rate at which vectors died, nor vector EIP. Host viremia, too, (alone and in interaction with CORT treatment) did not affect either rate. Null results are always difficult to interpret, but our study design provides some insight. First, it is unlikely that null results are driven by sample size; although we had fairly few birds on which to feed mosquitoes, we tracked survival and EIP in a large number of mosquitoes. It is thus unlikely that non-significant effects of CORT were due to low statistical power. Second, we revealed >75% of mosquitoes became infectious by 12-days post-feeding. This duration is consistent with other studies, but ours is the first (of which we are aware) that has estimated infection rates using passerines as hosts.

Our third discovery is perhaps the most enlightening ecologically; we discovered that even control-implanted finches, which never reached 10^5^ pfu ml^−1^ virus in circulation, were comparably infectious to *Culex* vectors as CORT birds. In our previous work, we conservatively recognized 5 logs as the transmission threshold birds must surpass to become infectious to vectors (Turell et al. 2000), however, others emphasize 4 logs (Kilpatrick et al. 2007; Tolsá et al. 2018). Here, it was clear that finches with a 4 log WNV titer, on average, can infect a key vector. However, we detected no effects of individual-level viremia on vector EIP or mortality rate in CORT-treated or control finches.

### Epidemiological implications of our data

We found no evidence that our CORT treatments altered the rate at which *C. quinquefasciatus* became infectious with WNV, nor did we observe any influence of CORT on vector mortality rates. These results indicate that any effects of host stress, as captured by our experimental approach, will manifest via other stages of the host-vector-virus interaction. Indeed, our other work clearly implicates host stress (as represented by sustained CORT elevations) as a potentially important driver of spatiotemporal variation in WNV risk. Identical CORT treatments to the ones used here make finches 2x more attractive to vectors than controls (Gervasi et al. 2016) and double the period of infectiousness (Gervasi et al. 2017a). The next step would be to integrate all of these effects into a single mathematical framework (Bergsman et al. 2016). We conducted a similar exercise recently with regard to light pollution effects on WNV viremia in house sparrows (*Passer domesticus*). Exposure to one form of light pollution, artificial light at night, extended the WNV infectious period of this avian species by 20% (Kernbach et al. 2019). When we estimated how this effect on host competence would change R_0_, the number of new infections expected to be generated by one infectious host in a wholly susceptible population, we found that risk increased by 41%.

Even though our data do not implicate host CORT as an important driver of EIP or mosquito viability, we hope that they inspire additional efforts to investigate the effects of anthropogenic stressors on emerging infectious diseases and especially zoonoses. Of course, many natural stressors such as predation risk, competition, co-infection, and other forces can have sublethal influences on hosts, which can alter the rates at which diseases emerge and spread (Buck et al. 2018; Mierzejewski et al. 2019). We encourage that special attention be directed to anthropogenic stressors such as pollutants, non-native species introductions, habitat loss and degradation, climate change, urbanization, and other human activities (Martin and Boruta 2014; Martin et al. 2010), as these are apt to alter host, vector, and parasite biology in diverse ways, none of which will have been common in the evolutionary history of most species. We also encourage additional work to reveal whether blood or protein composition is affected in a manner that might impact mosquito productivity or EIP, in spite of the null effects we detected with our experimental design. Finally, we encourage research of stress hormone effects on other avian and vector species, as competence varies extensively both within and among host *and* vector species (Tolsá et al. 2018; Turell et al. 2005). Some species might be more susceptible than others to stressors (Martin et al. 2010; Paull et al. 2011), and further, vector and host behaviors and densities might change contingent on context (Levine et al. 2013) (Apperson et al. 2004; Goodman et al. 2018). Even some individuals might contribute more to local epidemics than others (Scott et al. 1990), highlighting the need to consider ecological context (Gervasi et al. 2015) as well as evolutionary history (Downs et al. 2019) when aspiring to manage disease risk (Martin et al. 2019a).

## Acknowledgements

We recognize NSF-IOS grant 1257773 and the USF College of Public Health for support.

